# Domain-specific embeddings uncover latent genetics knowledge

**DOI:** 10.1101/2025.03.17.643817

**Authors:** S. S. Ho, R. E. Mills

**Affiliations:** Department of Human Genetics, University of Michigan, Ann Arbor, MI, USA; Department of Computational Medicine and Bioinformatics, University of Michigan, Ann Arbor, MI, USA

## Abstract

The inundating rate of scientific publishing means every researcher will miss new discoveries from overwhelming saturation. To address this limitation, we employ natural language processing to overcome human limitations in reading, curation, and knowledge synthesis, with domain-specific applications to genetics and genomics. We construct a corpus of 3.5 million normalized genetics and genomics abstracts and implement both semantic and network-based embedding models. Our methods not only capture broad biological concepts and relationships but also predict complex phenomena such as gene expression. Through a rigorous temporal validation framework, we demonstrate that our embeddings successfully predict gene-disease associations, cancer driver genes, and experimentally-verified protein interactions years before their formal documentation in literature. Additionally, our embeddings successfully predict experimentally verified gene-gene interactions absent from the literature. These findings demonstrate that substantial undiscovered knowledge exists within the collective scientific literature and that computational approaches can accelerate biological discovery by identifying hidden connections across the fragmented landscape of scientific publishing.

## Introduction

The sheer saturation and overwhelming growth of scientific literature means that every researcher will miss the dissemination of new datasets, discoveries, and knowledge. One way to overcome human limitations in text synthesis is by employing machine-learning methods formulated for language, known as natural language processing (NLP)^1^. NLP has been utilized across numerous fields to perform tasks such as sentiment analysis^2^, recommender systems^3^, information retrieval^4^, and machine translation^5^. Specifically, NLP has been used in the life sciences to automate literature curation^6^ and gene identification^7^, and has become a targeted domain for the development of pretrained large language models (LLMs)^8–11^. Notably, most of these applications have focused on automated comprehension and generative capabilities with little evaluation of the ability to derive latent insights.

Extracting latent knowledge from language models presents a fundamental challenge: how do we assess models on information they have not explicitly learned? Seminal work from Tshitoyan et al.^12^ address this challenge through temporal validation: training models exclusively on text published up to a specific year and evaluating the model’s abilities to predict relationships that appear in literature years later. This approach produces a clear delineation between what a model has seen and what a model does “not know,” which represents novel information.

While LLMs represent state-of-the-art in NLP, they require extensive pre-training processes with hundreds of gigabytes of text data. This substantial data requirement obscures the boundaries between known and unknown information, making it challenging to identify what these models do ‘not know.’ Thus, their dependence on massive, heterogeneous datasets coupled with expensive training costs makes them unsuitable for temporal validation methods.

We instead utilize embedding methods, which aim to encode text as semantically-informed numerical vectors. Embedding approaches offer crucial advantages: manageable computational demands, unified representations that simplify downstream analysis, and critically, precise control over training corpora to enable a temporal validation framework. Recent studies have demonstrated that these methods can effectively extract latent knowledge in fields such as pharmacology^13^, materials science^14^, and electrochemical engineering^15^. Such approaches are thus likely able to predict previously unidentified relationships in genetics and genomics literature.

Here, we construct a corpus consisting of over 3.5 million abstracts coupled with a custom processing pipeline to robustly capture semantic knowledge through various embedding techniques. Our results show that these embeddings capture broad biological trends but also encode surprising amounts of biological information within their representations. Building upon these findings, we advance beyond prior studies that primarily utilized dot product similarities between input and output embeddings^12–15^ by implementing downstream machine-learning models that directly leverage the embeddings and their interactions. We demonstrate the ability of our embeddings and models to reveal latent knowledge, successfully predicting interactions that are validated in experimental data but never mentioned in the literature.

## Results

### Gene and disease normalized scientific corpora

Word embeddings encode the semantic context of a given vocabulary instance: it is imperative that the final corpus used for model training is cleaned and normalized to optimize the fidelity of learned representations. First, we must reduce instances where the same word can mean semantically different things, such as the definition of “promoter” in the context of the genetics versus the context of business. Second, we need to minimize instances of vocabulary redundancy where different notations refer to the same entity, such as “P53” and “TP53”, which fragments the learned semantics into separate vector representations. To address field-specific definitions, we trained multiple classifiers on term frequency-inverse document frequency using a fine-tuning approach. Our classifier achieved an F1 score of 93% via five-fold cross validation (Supplementary Table S1; see Methods), ensuring our corpus maintains high relevance to genetics and genomics. To address word normalization, we apply the biomedical normalization tool HunFlair2^16^ to normalize genes and disease entities. Our pre-processing pipeline resulted in a custom corpus of ∼3.5 million genetics and genomics relevant, normalized abstracts optimized for model training (Fig 1).

**Fig 1.**
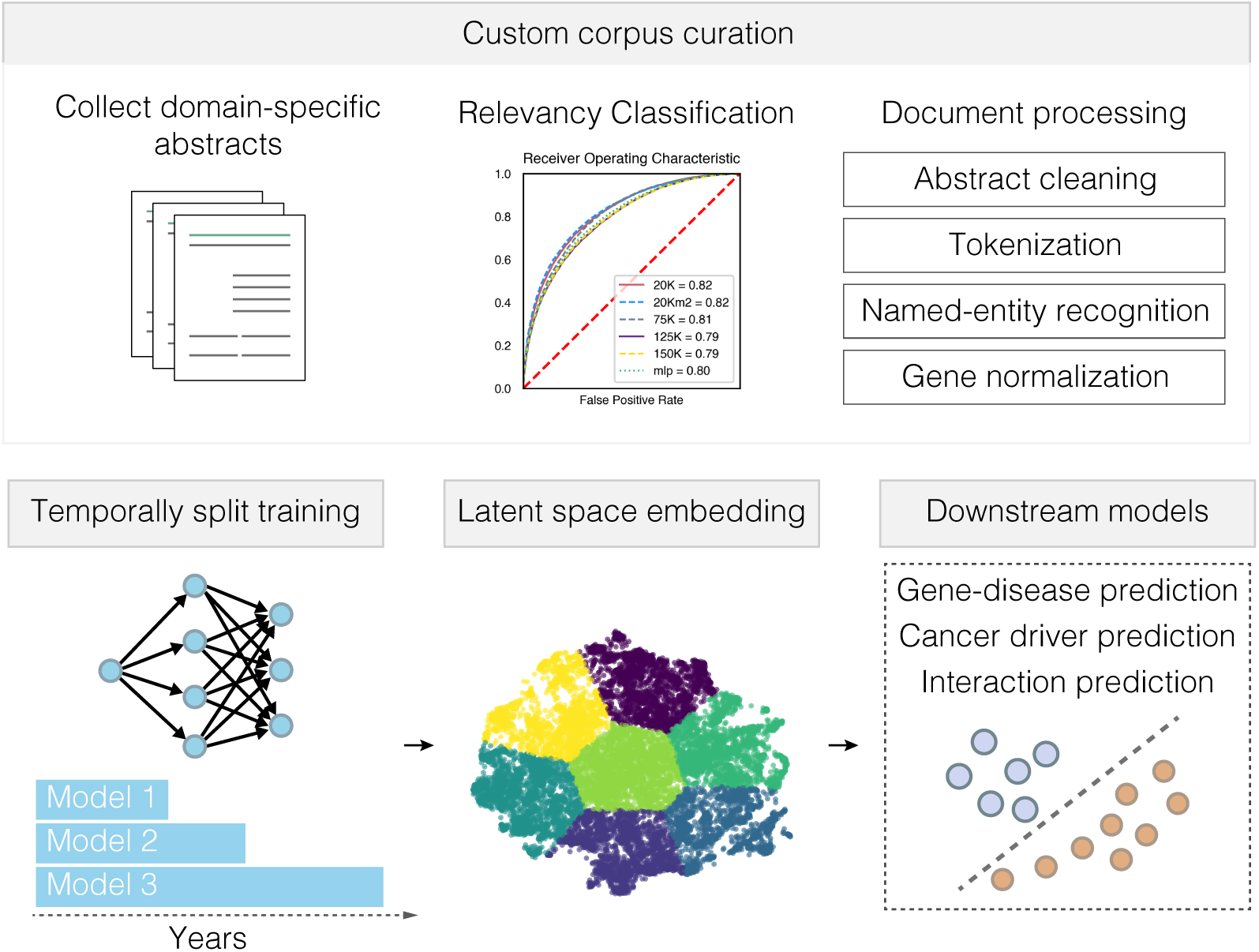
Natural language processing workflow. Our pipeline begins with curation and processing of our custom corpus in a temporally-aware manner. We build embeddings using complementary approaches and use them as input into downstream models designed to evaluate for the presence of latent knowledge.

We additionally extract gene-gene and gene-disease co-occurrence networks from our corpus to capture information beyond semantic encodings. For each abstract, we track co-occurrence using the normalized entities along with database aliases (see Methods). This process yields gene-gene and gene-disease co-occurrence graphs that serve as input for our embedding models. We employ two complementary embedding approaches: word2vec^17^, which learns vector representations by predicting neighboring words in text, and node2vec^18^, which optimizes embeddings based on network traversal. Together, these methods capture both contextual semantic relationships and co-occurrence patterns, providing a comprehensive representation of biological relationships encoded in literature.

Word embeddings recapitulate domain-specific knowledge

To verify the implementation of our custom processing pipeline (Fig. 1) and the quality of our representations, we first analyzed the ability of trained embeddings to recapitulate domain-specific knowledge. We referred to the classical embedding analogies posited by Mikolov et al.^17^ which demonstrate that vector operations reflect semantic relationships, such as the famous example:

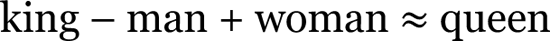

Our embedding models exhibit similar but domain-specific capabilities. For instance, our models capture broad, biologically relevant analogies, such as:

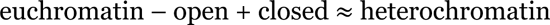

The captured analogies span more than broad generalizations. We show that the embeddings capture complex relationships such as those capturing gene-disease associations (cystic fibrosis – *CFTR* + *SMN1* ≈ spinal muscular atrophy) and pathway-specific interactions (*PIK3CA* – *AKT1* + *RAF1* ≈ *KRAS*). We projected these embeddings and their relationships onto two dimensions using principal component analysis (Fig. 2a,b,c) and find that analogies exhibit similar directions and patterns in the vector space, suggesting that the embeddings encode biological concepts.

**Fig 2.**
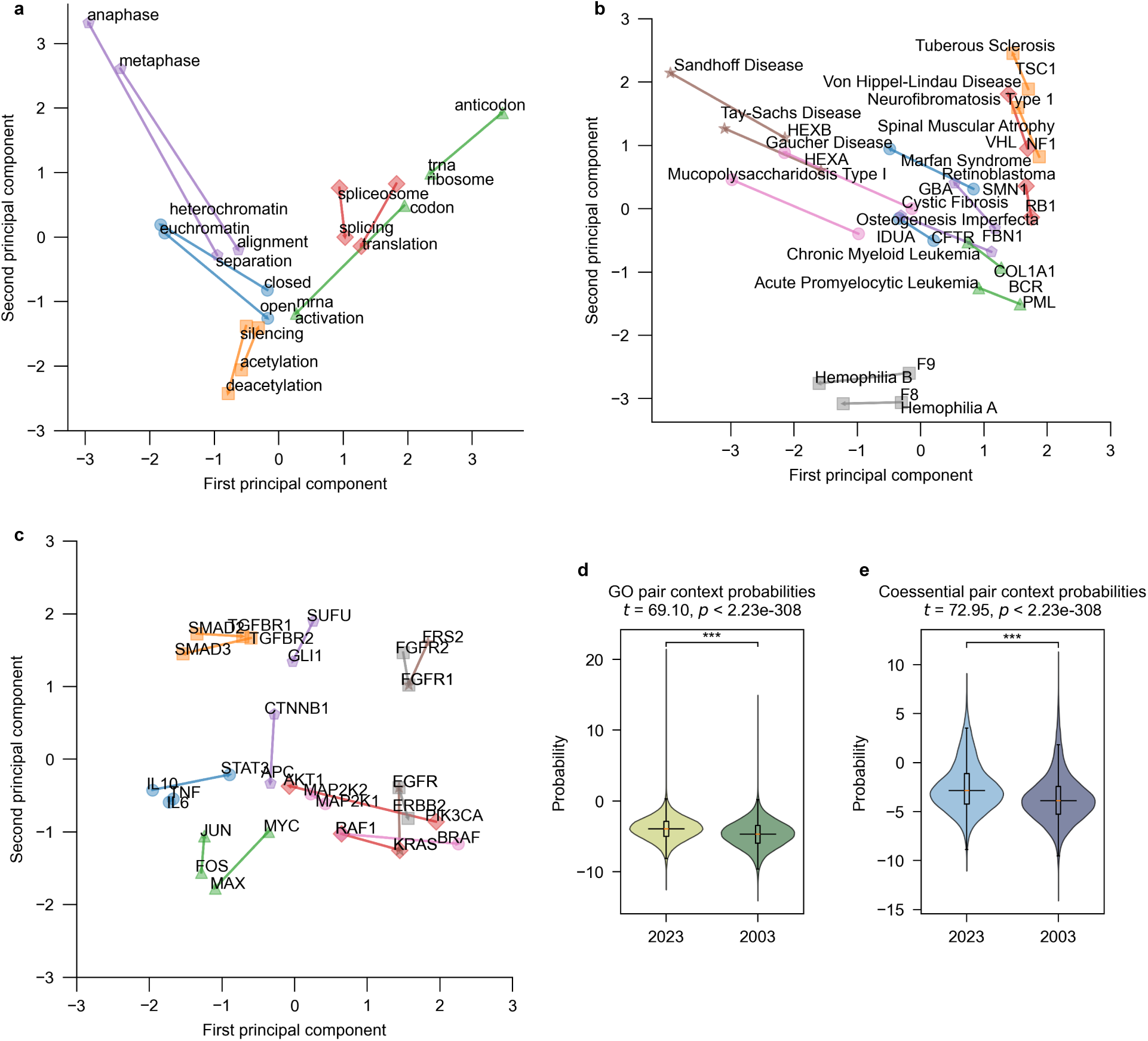
Embeddings encode known biological relationships. a, The semantics embedded into vectors recapitulate broad scientific relationships through by vector operations that look for similarities in Euclidian space. The similar direction and magnitude of these vectors suggests that the embeddings capture meaningful relationships. For example, as “anaphase” is characterized by “separation”, “metaphase” is characterized by “alignment”. The depicted relations are of broad biological analogies, b, known gene-gene interactions, and c, gene-disease associations. d, Co-occurrence predictions between 10,000 randomly sampled gene-gene interactions that overlap GO pairs and. e, gene co-essentiality.

Next, we investigated whether our embeddings could recapitulate known biological relationships between genes in a quantifiable manner. We used the dot product between the hidden and output layer embeddings to measure co-occurrence probability between individual units of text (tokens). We assessed these probabilities on two ground truth datasets: gene pairs sharing Gene Ontology (GO)^19^ processes and pairs from an atlas of gene co-essentiality^20^. We compared embedding models trained on abstracts up to 2003 to those trained on abstracts through 2023, revealing a significant difference in the co-occurrence probabilities between the two time periods for both the GO pairs (t = 69.10, *P* < 2.23e-308; Fig. 2d) and co-essential pairs (t = 72.95, *P* < 2.23e-308; Fig. 2e). Models trained on the entirety of the corpus showed higher co-occurrence probabilities for biologically related pairs than those trained on the earlier corpus suggesting the embeddings are capturing the accumulation of biological knowledge in scientific literature as discoveries emerge.

### Embeddings accurately predict gene expression

Building on these analyses, we sought to evaluate whether our embeddings could predict biological properties as complex as gene expression. Using our full corpus (2023), we used embeddings to train a gradient boosted model to predict expression levels from the GTEx v8 dataset^21^. The model achieves a strong correlation (Pearson *r* = 0.6016) on hold-out test genes (Fig. 3a), outperforming GenePT^22^, a transformer-based embedding method of larger dimensionality (Supplementary Fig. 1) and significantly outperforms the random baseline model trained with Xavier uniform^23^ embeddings (Fig. 3b). These results demonstrate that our embedding approach not only captures qualitative semantic relationships but also encodes quantitative biological properties that can be leveraged for predictive modelling of complex biological patterns like gene expression.

**Fig 3.**
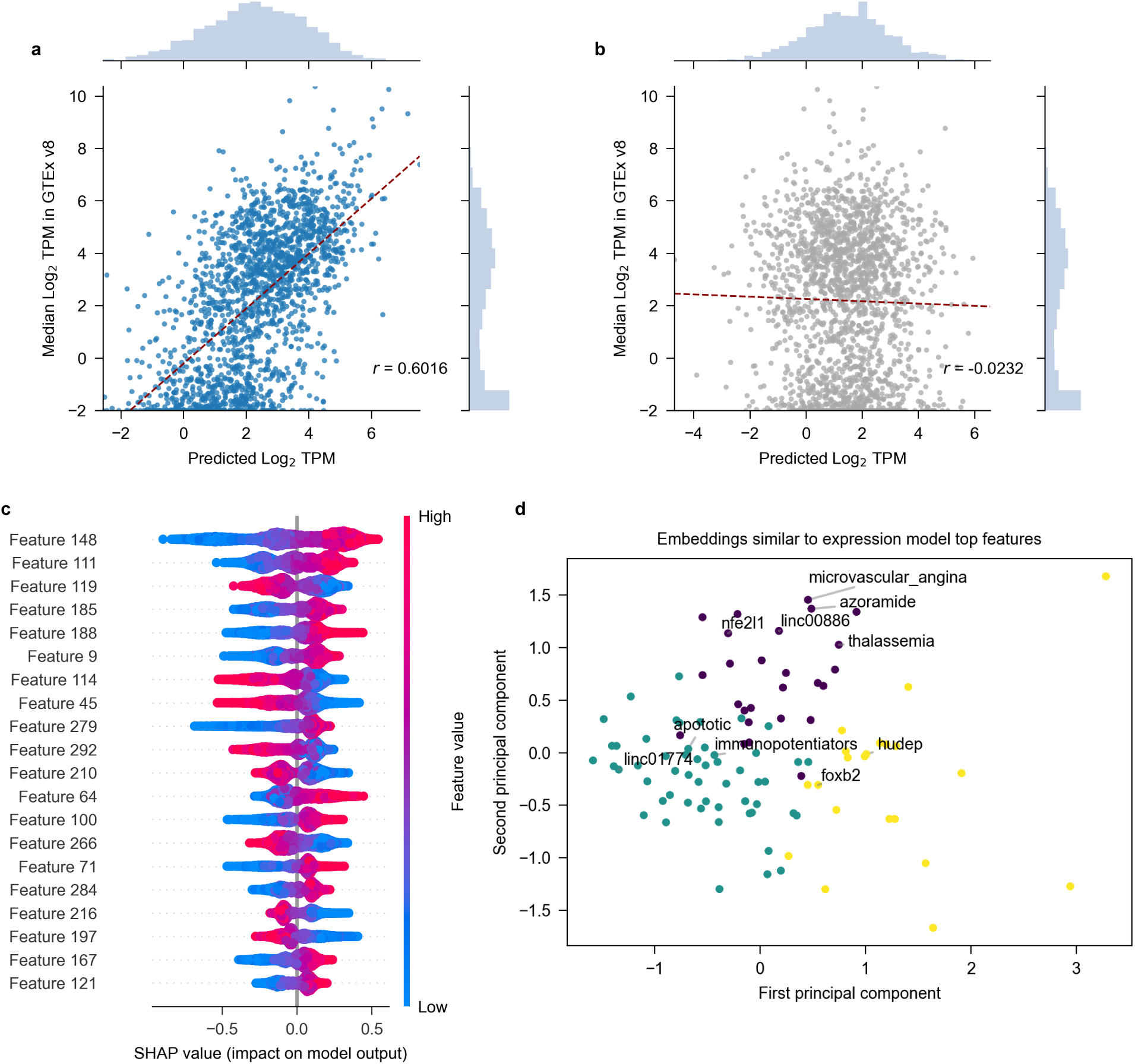
Word embeddings accurately predict gene expression. a, Scatter plots of predicted versus expected gene expression for hold-out test genes (chr 1) with best fit line indicated in red. Model trained with word2vec embeddings achieves a Pearson r = 0.6016 predicting the median TPM of genes from the GTEx v8 dataset. b, expression prediction performance using random embeddings populated with Xavier initialization (Pearson *r* = −0.0232). c, SHAP values for the top 20 features in the gene expression prediction model, showing the impact of each embedding dimension on model output. Red areas indicate where higher feature values increase predicted expression, while blue areas show where higher feature values decrease predicted expression. d, Principal component analysis visualization of words with embeddings most similar to high-SHAP-value features, revealing diverse biological terms spanning multiple domains including diseases, transcription factors, cellular processes, and non-coding RNAs.

To understand which embedding dimensions contribute the most to expression prediction, we applied SHAP (Shapley Additive exPlanations)^24^ analysis to identify the most influential features (Fig. 3c). We identified the top 100 embeddings most similar to these high SHAP-value features and projected them onto two dimensions using principal component analysis (Fig. 3d). The semantic space associated with the influential embedding dimensions revealed disease terms (“microvascular_angina”, “thalassemia”), transcription factors (“foxb2”), cellular processes (“apoptotic”), drug-related terms (“azoramide”, “immunopotentiators”), and long-noncoding RNAs (“linc00886”, “linc01774”). This suggests that the embedding dimensions deemed important by the model do not map cleanly to single biological concepts or relationship types but rather encodes multifaceted information that spans multiple biological domains. The blending of signals within these embedding dimensions highlights how semantic knowledge can exist distributed through seemingly disparate text concepts.

### Predicting disease-causative genes before their identification in literature

To test explicit latent knowledge extraction, we next investigated whether our trained embeddings could predict biological phenomena before their documentation in literature. We collected temporal provenance data tracking the year genes were identified as disease causal^25^. Using this chronology, we constructed a series of models using a temporal validation framework, training models with cut-off years to limit their training data. For example, for the 2003 model, we trained the model on gene-disease co-occurrences on a corpus up to 2003. We then tested the model on the collected causative gene-disease relationships discovered after 2003, after the model’s training cutoff year. This experimental design creates a rigorous temporal validation framework allowing us to assess whether the embeddings identify causative gene-disease relationships before their formal characterization in the literature.

Our framework and validation experiments demonstrate that embedding-based models can effectively predict future causative gene-disease relationship, consistently outperforming random baselines across all time periods test (Fig. 4a; Supplementary Fig. 2). Notably, models utilizing word2vec and node2vec embeddings showed comparable performance. All models achieved average precision (AP) values > 0.60, with the most performant model achieving an AP = 0.7411. At a probability threshold of 0.75, our best performing model (XGBoost; 2009) successfully identified 51 out of 74 gene-disease associations (68.92% accuracy) that are confirmed in the literature after the training cut-off. We emphasize that the performance of these models must be contextualized with the size of the test set (Supplementary Fig. 3): as the temporal cut-off advances, more genes-disease relationships are discovered, and the size of the test set decreases.

**Fig 4.**
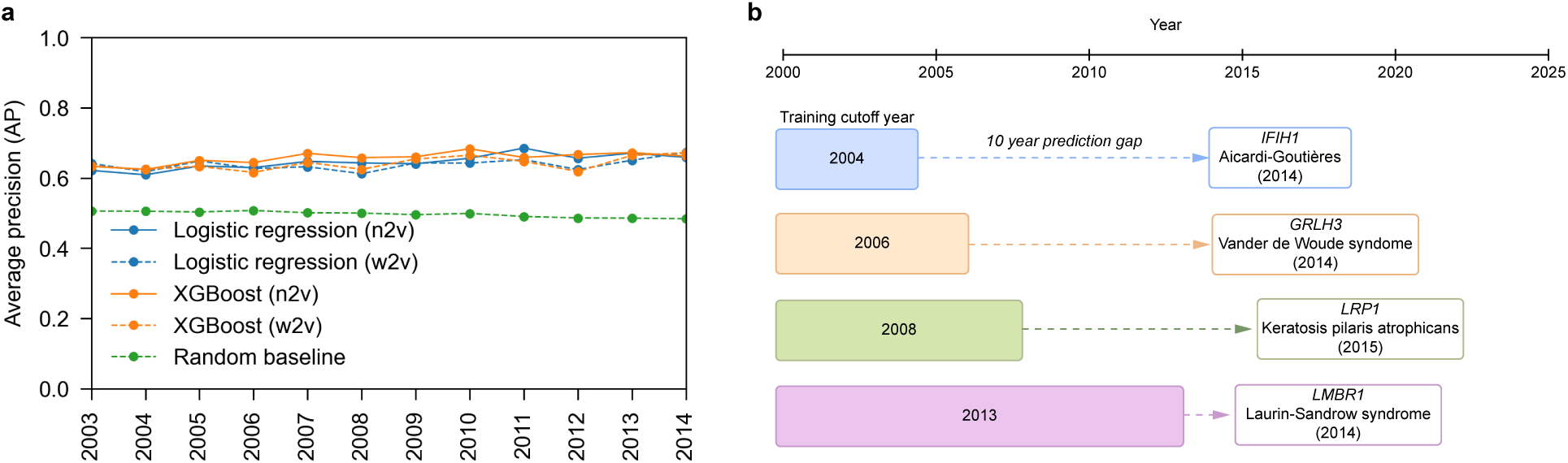
Predicting gene-disease associations before their appearance in literature. a, Average precision of models trained over the temporal split. Each model is trained on the gene-disease co-occurrences up to its training year and tested on gene-disease provenance data for years after the corpus cutoff. b, Examples of time gaps between model predictions and years that genes are described as causal drivers of different diseases ranging from 1 year to 10 years.

We inspected some of the model predictions after training. For example, a model trained on a corpus up to 2004 predicted a high probability (0.925) of association between Aicardi-Goutières syndrome and *IFIH1*, which was identified as causative in 2014^26^, ten years after the training cut-off of this corpus (Fig. 4b); we confirmed that the two terms never co-occur in our 2004 corpus. We find similar patterns across other models: a 2006 model predicts an association between *GRHL3* and Van der Woude syndrome (0.988) which was discovered 8 years later^27^, a 2008 model predicts high probability between *LRP1* and keratosis pilaris atrophicans for which a causative mutation was discovered 7 years later^28^, and a 2013 model predicts *LMBR1* as likely associated (0.774) with Laurin-Sandrow Syndrome, formally discovered the next year, in 2014^29^ (Fig. 4b). These examples demonstrate that our embeddings capture pertinent signals and validate our framework’s effectiveness in evaluating the capacity to extract latent knowledge from scientific literature.

### Predicting cancer drivers before their identification in the literature

We continued testing latent knowledge extraction by collecting temporal provenance data tracking the year genes were first identified as putative cancer drivers^30,31^. We constructed a series of temporally-split models trained on embeddings from the beginning of our corpus to years 2001 through 2018 where cancer driver genes discovered up to the training year were used as training examples and cancer drivers discovered after the cutoff were used as test examples.

In line with our results from gene-disease associations, our temporal validation experiments demonstrated that embeddings effectively predict future cancer driver gene discoveries and outperform random baselines across all time periods tested (Fig. 5a). Precision-recall curves show all embedding approaches maintained substantial advantage over the random baselines (Supplementary Fig. 4).

**Fig 5.**
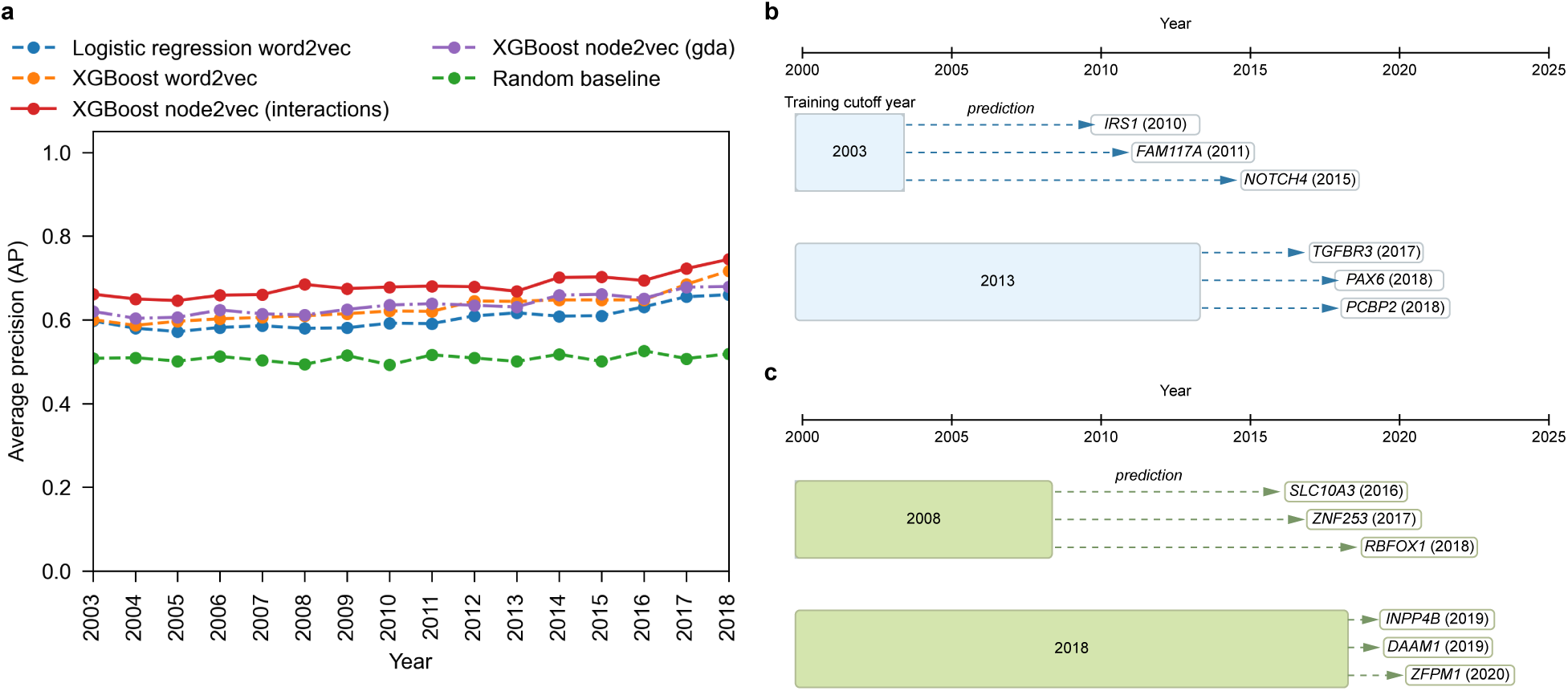
Predicting putative cancer drivers before documentation of their driving mechanism. a, Average precision metrics for cancer-driver gene prediction models trained up to specific cut-off years. The gradient-boosted model leveraging node2vec with gene-gene co-occurrence data consistently outperforms other approaches across all temporal splits maintaining average precision values between 0.65-0.75 throughout the evaluation period. b, Identification and validation of cancer-driver genes using a temporal validation framework. Word2vec trained up to a year cut-off (x-axis) predict cancer driver genes before their driver mutations are identified in the literature. c, Node2vec models.

Notably, the XGBoost model leveraging node2vec with interaction data demonstrated superior performance, maintaining AP values between 0.65-0.75 throughout the evaluation period with a modest upward trend in later years. This model achieved higher average precision (PRAUC=0.74; 2018) compared to the word2vec model (PRAUC=0.71; 2018), suggesting that the additional network structure information enhances cancer driver gene identification. We noticed that stability started to degrade as the temporal split progressed (Supplementary Fig. 4) and reasoned that this degradation stems from the increasing number of known cancer driver genes over time, which consequently reduces the test set size (Supplementary Fig. 5). Re-evaluation of the models using a limited three-year prediction horizon demonstrated continued robust performance (Supplementary Fig. 5), indicating that embeddings capture generalizable signatures of cancer driver genes over time-specific patterns.

We examined specific examples of successful predictions from our temporal validation models. For instance, our word2vec-based model trained on data up to 2003 assigned high confidence scores to *IRS1*, *FAM117A*, and *NOTCH4*, which were formally identified as cancer driver genes in 2010^32^, 2011^33^, and 2015^34^, respectively (Fig. 5b). Later models demonstrated similar predictive power, with a 2008 model identifying *SLC10A3* (2016)^35^, a 2013 model identifying *TGFBR3* (2017)^36^, and a 2018 model identifying *INPP4B* and *ZFPM1* which were formally recognized as cancer drivers in 2019^37^ and 2020^38^, respectively (Fig. 5b,c). More examples are illustrated in Fig. 5.

It is pertinent to note that false positives inevitably exist in our models. Our test set does not include all cancer-related genes but rather putative drivers. This means that some cancer-gene associations are encoded in the embedding space before their driving mechanism is explicitly documented, allowing our models to leverage these latent semantic relationships to prioritize potential cancer drivers. However, we argue that this is analogous to how researchers draw connections from partial data to invest in promising avenues of inquiry. In this sense, although NLP is not perfect, it accelerates the exploratory process by rapidly identifying and exploiting these underlying relationships despite the presence of false positives.

### Predicting experimentally-validated interactions from text representations

We then investigated whether our embedding models could predict experimentally verified gene-gene or protein-protein interactions. This task presents a fundamentally different challenge: whereas gene-disease associations and cancer driver genes are explicitly discussed in literature, most gene-gene and protein-protein interactions exist primarily as experimental data. This makes interaction prediction inherently more difficult as the model must infer relationships with minimal to no text evidence. We used gene co-occurrence in abstracts as positive training examples and evaluated models on 165,689 experimentally-validated interactions compiled from a gene co-essentiality map^39^, physical protein-protein interactions^40,41^, and single-cell gene co-expression data^42^, thus simulating the real-world scenario of predicting novel interactions from existing literature. Our temporal validation experiments demonstrate that embeddings consistently predict interactions that are never documented in literature: XGBoost models trained on word2vec embeddings maintained average precision values consistently exceeding 0.80 throughout the entire evaluation period, consistently outperforming random baselines (Fig. 6).

**Fig 6.**
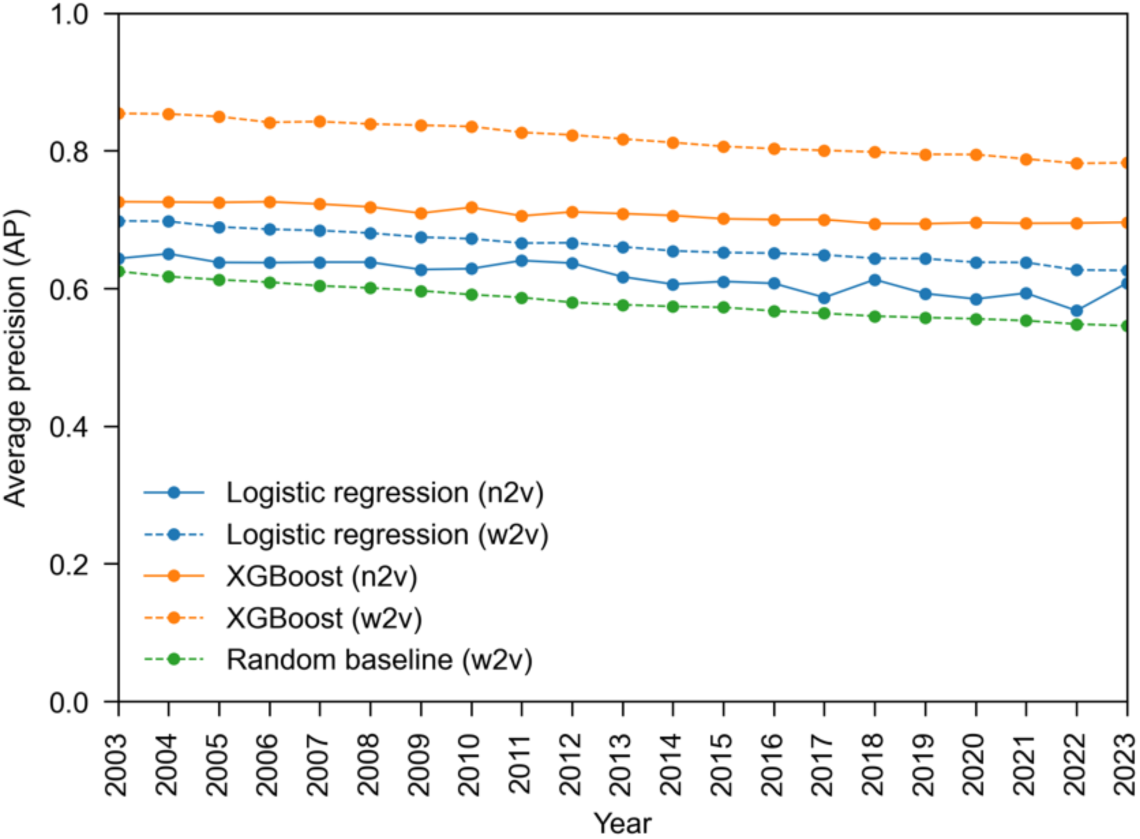
Predicting interactions that remain undocumented in literature. Average precision (PR AUC) for interaction prediction models spanning a 21-year temporal split. Gradient boosted models trained on word2vec embeddings (orange, dashed lines) consistently outperform all other configurations.

The pronounced performance gap between word2vec and node2vec embeddings for interaction prediction (Fig. 6; Supplementary Fig. 6) presents an interesting contrast to our cancer driver prediction results, where node2vec demonstrated slight advantages, and to our gene-disease analysis, where methods performed comparably. This difference suggests that the optimal embedding approach may depend on the specific prediction task: word2vec’s direct capture of contextual patterns appears particularly suited for relationship prediction, while node2vec’s incorporation of network structure may better group genes with properties relevant to cancer drivers.

We examined successful model predictions and found an interaction between two breast cancer related genes, *PTGES3* and *MCM7*, with a high probability score (0.929). We find these two genes are never co-mentioned in the same abstract and further find that this interaction does not exist in StringDB (v.12.0)^43^, a widely used protein-protein interaction database. The interaction was documented in BioGrid^44^ with a singular source of evidence: experimentally validated physical contact from the OpenCell compendium^41^. Similarly, our model predicted an interaction between *ACVR1B* and *HSP90AB1* (0.932) that does not appear in either StringDB or BioGrid but exists solely as a verified binary interaction in the Rolland et al. dataset^45^. These examples demonstrate that our embeddings successfully infer experimentally-validated relationships even when direct textual evidence of the interaction is entirely absent.

### Static embeddings out-perform transformer-derived embeddings

Our work makes extensive use of static embeddings due to enhanced control over corpora and computational efficiency. While static embeddings are performant with the correct problem framing^46^, methods that employ contextual embeddings, such as ELMo^47^, BERT^48^, and GLUE^49^, are considered state-of-the-art. We sought to further evaluate our embeddings through comparisons against a fine-tuned bidirectional transformer model based on BiomedBERT^50^. We augment the model with our normalized entities, fine-tune on our corpus using masked language modelling, and derived embeddings by either averaging representation occurrences or using an attention pooling mechanism (see Methods). As we cannot control for the pre-training corpus, we compare these transformer-derived embeddings and our static embeddings using five-fold cross-validation for the gene-disease and cancer driver models, while maintaining the standard train/test split for our interaction model.

Performance comparison across our three prediction tasks revealed mixed results. The transformer-derived embeddings show improvements for gene-disease association prediction (AUC: 0.9911 vs. 0.9731), but our static embeddings demonstrate superior performance for cancer driver prediction (AP: 0.7672 vs. 0.7734) and for interaction prediction (AUC: 0.7579 vs 0.7827). While the transformer model achieved better results in one task, these improvements came at significantly higher computational expense at every step of their implementation: at fine-tuning, embedding extraction, and downstream model training. These results demonstrate that carefully constructed domain-specific embeddings can achieve similar performance to, and even exceed, computationally intensive contextual models while maintaining practical feasibility.

## Discussion

We have demonstrated that domain-specific embeddings trained on carefully processed corpora can effectively predict gene-disease associations and cancer driver genes years before their formal documentation and identify experimentally verified gene interactions absent from literature. The complementary embedding approaches showed task-specific strengths, highlighting that knowledge in scientific literature exists in multiple dimensions as contextual semantics and network structures derived from co-occurrence patterns. Our results demonstrate that substantial undiscovered knowledge exists within the collective scientific literature and can be computationally extracted. Additionally, we demonstrate the importance of input data preprocessing and normalization, out-performing state-of-the-art transformer-based embeddings with a fraction of the computational cost. While NLP approaches have been applied to biomedical corpora, to our knowledge, none have employed an extensive validation framework to assess latent knowledge extraction.

Challenges in biomedical NLP remain. Two major obstacles we faced were the lack of field-standardized vocabulary and insufficient provenance data. Without proper normalization, semantically identical contexts become fragmented across multiple representations, diluting embedded signals. For example, the HGCN alias set contains 147,237 ways to mention 42,861 genes, while the CTD Diseases set includes 74,802 ways to mention 13,307 different diseases. Disease terminology exhibits particularly severe semantic fragmentation compared to gene terminology where diseases appear under numerous permutations while genes like “CTCF” have relatively few linguistic variations. Additionally, biology severely lacks the temporal metadata required for validation frameworks, with few publicly available and thoroughly vetted provenance datasets, limiting the scope of hidden knowledge validation.

Beyond these technical challenges, knowledge extraction from scientific literature is influenced by the current structure of scientific communication. The academic publishing ecosystem, while valuable for disseminating research, prioritizes broad applicability over detailed mechanistic descriptions. This manifests in two notable ways: first, a degree centrality issue where certain genes and diseases receive disproportionate focus compared to potentially significant but less studied biological entities; second, the scholarly incentive to demonstrate broader relevance by referencing numerous entities introduces false positives into text-based analyses. Moving forward, balancing specialized mechanistic descriptions with demonstrations of broader relevance would significantly improve the task of synthesizing biological knowledge from natural language.

In summary, our work contributes a rigorous framework for latent knowledge extraction from biological text and demonstrates how these methods can generate guided hypotheses about gene functions and interactions. This approach complements traditional experimental methods and offers a powerful approach for navigating the increasingly complex written landscape of genomic knowledge.

## Author contributions

S.S.H. conceived the projected, curated the data, annotated the abstracts, trained the models, performed all analyses, and wrote the manuscript. R.E.M. supervised the project.

## Data and materials availability

All code is available at https://github.com/sciencesteveho/genomic_nlp.

## Acknowledgements

We thank U. Kadiyala, A. Weber, W. Zhou, P. Ghandi, C. Sun, W. Gu, Y. Wang, and M. Sherman for help with downloading data.

## Methods

### Data collection and relevance classification

Approximately ∼8.6 million (8,585,476) abstracts related to genetics, genomics, and molecular biology were collected from the SCOPUS API using pybliometrics (v.3.6)^51^. The resulting literature were published between 1989 to 2023 relating to 123 genetics and genomics relevant search terms (Supplementary Table 2). Only entries with types “article”, “review”, or “letter” were kept. The titles of document were appended to the beginning of each abstract so that they would be used as training data. Abstracts were cleaned of copyright, journal, and publishing tags with regular expressions.

### Relevancy classification

Previous work has shown that embeddings perform best when relevant texts are curated as opposed to maximizing corpus size^12^. We manually annotated 2,014 randomly selected abstracts as “relevant” or “not relevant” for text classification based on the following:

1. The abstract must mention at least one molecular feature at the DNA, RNA, or protein levels.
2. The abstract should specifically describe relevancy of the molecular feature to a DNA, RNA or protein product, a molecular mechanism, or a human disease.
3. The abstract should be relevant to mammalian mechanisms. Abstracts specifically targeted at other eukaryotes, such as plant cultivars, bacteria, or fungi considered “not relevant.”

1,026 abstracts were annotated as “relevant” while 988 abstracts were annotated as “not relevant”. We trained multiple logistic regression and MLP classifiers varying term frequency-inverse document frequency (tf-idf). Classifiers were first trained on trained on 40,000 abstracts stratified by journal type as a measure for relevancy (Supplementary Table 3). We then fine-tune the model on the smaller set of manually annotated abstracts with adjusted hyperparameters to avoid overfitting and evaluate all models with five-fold cross validation. The cleaned corpus consists of 3,459,487 relevant abstracts.

### Named entity normalization and abstract processing

We used HunFlair2 (v.0.14.0) for named entity recognition (NER) and named entity normalization (NEN) in our dataset to ensure the robust capture of semantics^16^. For example, if the gene “TP53” exists as tokens “TP53” and “TP53+” the semantic information would effectively fragment across embeddings. NEN standardizes genes across different contexts, normalizing instances of “TP53+” to a shared “TP53”. We apply HunFlair2’s gene and disease linkers to normalize gene and disease tokens, respectively.

Abstracts were tokenized using the “en_core_sci_lg” model from the biomedical NLP package scispaCy (v.0.5.4)^52^. We casefolded all tokens, removed special characters, punctuation marks, and replaced standalone numbers with a universal symbol “<nUm>”. We generated *n*-grams (phrases) with strict thresholds, requiring phrases to appear a minimum of 75 times and with a score of 60. We repeated phrase generation four times, generating a maximum of 8-gram phrases. Genes and diseases were excluded from *n*-gram generation to avoid dilution of our normalized entities.

Abstracts were annotated with the year they were published and allowed us to produce a temporal split. We built individual corpora from 2003 to 2023, where each year signifies publication cut-off year. This allowed us to train models and strictly evaluate their ability to predict information from future abstracts beyond the range of their training data.

### Co-occurrence graph extraction

We extracted entity co-occurrence graphs based on either the presence of genes with genes or genes with diseases. First, we mapped each gene to its possible aliases and each disease to its possible aliases as listed in the HUGO Gene Nomenclature Committee and Comparative Toxicogenomics databases, respectively, and added the HunFlair2 normalized name to the alias mapping. Then, for each abstract we tracked the co-occurrence of genes and diseases via the presence of the normalized name or any aliases. We produced gene-gene co-occurrence graphs and gene-disease co-occurrence graphs for each of the temporally split corpora.

### Embedding models

We trained embedding models based on word2vec for tokenized texts and node2vec for the co-occurrence graphs. For comparison of the gene-expression prediction task, we downloaded 1,536-dimensional embeddings from GenePT^22^ and used their embeddings trained via ChatGPT’s “text-embedding-ada-002’ model.

We trained word2Vec embedding models using Gensim (v.4.3.3)^53^ on the 8-gram transformed corpora with the following parameters: 300-dimensional embeddings, minimum word count of 8, window size of 10, initial learning rate of 0.01 decreasing to 0.0001 in 30 epochs, a subsampling threshold of 0.0001, and 15 negative samples using the skip-gram algorithm.

We trained Node2vec embedding models on the gene-gene co-occurrence graphs using the GRAPE (v.0.2.4) and Embiggen graph representation packages^54^ with the following parameters: 128-dimensional embeddings, walk length of 128, window size of 5, and learning rate of 0.01 over 30 epochs.

### SHAP analysis

We implemented SHAP (SHapley Additive exPlanations) values for gradient-boosted decision tree models through the SHAP python package (v.0.46.0) to interpret feature importance of our embedding dimensions. For each model, we generated SHAP values using TreeExplainer on scaled input features from the training dataset. We identified the top contributing embedding dimensions by calculating the mean absolute SHAP values across samples and selecting the highest-ranked features. These key dimensions were used to filter gene embeddings based on cosine similarity to a reference vector derived from the training set. The filtered embeddings were then clustered using K-means (k=3) and visualized through principal component analysis to identify groups of embeddings with similar patterns in the most predictive embedding dimensions.

### Fine-tuned transformer model

We fine-tuned a bidirectional transformer model (BERT) using the Hugging Face Transformers library (v4.44.2). We chose ‘microsoft/BiomedNLP-BiomedBERT-base-uncased-abstract-fulltext’ as our base model for its pre-training on biomedical abstracts. We augmented the model’s vocabulary by adding custom tokenizer entries for our normalized gene and disease entities. The model was trained using a masked language modeling objected with 15% token masking probability for 3 epochs with an effective batch size of 128, a learning rate of 2e^-5^, bfloat16 mixed precision, AdamW optimizer with weight decay 0.01, and a linear learning rate scheduler with 10% of total training steps as warm-up, saving the best model according to the lowest training loss.

Following fine-tuning, we extracted average-pooled gene and disease entity embeddings by processing abstracts through the model and computing the mean of the final hidden layer representations for each entity token across all its occurrences in the corpus:

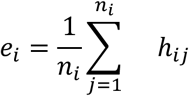

Where *e*_i_ is the final embedding for entity *i*, *n*_i_ is the number of occurrences of entity *i*, and *h*_ij_ is the hidden representation of the *j*th outcome.

We enhanced our embeddings using an attention mechanism that weights contextual representations by importance. A single-headed attention layer with a linear projection computes normalized importance scores α_i_ for each entity occurrence:

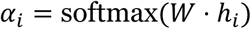

Where *h*_i_ represents the final hidden layer representation of the *i*^th^ occurrence and *W* is a learned weight matrix projecting the hidden dimension to a scalar score. The final attention-derived embedding for each entity *e* was computed as the weighted sum of hidden representations:

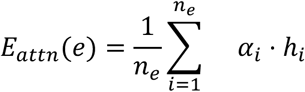

Where *n*_e_ is the number of occurrences of entity *e* across the corpus.

### Gene expression prediction

We used trained word embeddings to predict gene expression levels from the Genotype-Tissue Expression (GTEx) consortium dataset (v.8)^21^. For each gene, we calculated its median TPM value across all samples and tissues and applied a log_2_ transformation with a pseudocount of 0.25. We filtered the genes for protein-coding genes with available trained embeddings and reserved all genes on chromosome 1 (1,817 genes) for testing. Genes from all other chromosomes (14,760 genes) were used for training.

Expression values were predicted via the gradient-boosted ensemble decision tree model XGBoost (v.2.1.1)^55^. To optimize hyperparameters, we ran a grid search with the specified (Supplementary Table 4) and evaluated via five-fold cross validation. To train the random baseline, we generated embeddings of the same dimension and shape and populated each vector via Xavier initialization, a method commonly used to generate embeddings for neural networks due to its training stability^23^.

### Gene-disease association prediction

We collected data from Ehrhart et al.^25^ which links over 3,000 causative genes to over 4,000 monogenic disorders with provenance data and trained a series of machine learning models using a temporal validation framework. For example, if the underlying model was trained with a cutoff of 2013, the positive training set only included causative gene-disease associations up to 2013, and the test set included genes identified as causative gene-disease associations after 2013. We randomly sampled negative training examples as those that do not overlap the provenance data or the gene-disease co-occurrences and filter the test set for any overlapping pairs to avoid data leakage.

We trained XGBoost and logistic regression models comparing to a random baseline that predicts a random probability between 0 and 1. Models were trained on the concatenation of two vectors to represent an interaction.

### Cancer gene prediction

We collected data from the COSMIC Cancer Gene Census and Network of Cancer Gene databases^30,31^ which annotate cancer genes with the publication from which their cancer driving mechanisms is discovered. We used Biopython (v.184)^56^ to append each input with the year of their associated PubMed ID to generate the provenance data.

We trained a series of machine learning models using a temporal validation framework (described above). We randomly sampled negative training examples as genes did not overlap the COSMIC, NCG, or Intogen^57^ databases. We trained XGBoost and logistic regression models and implemented a random probability baseline (described above).

### Interaction prediction

We collected a gene co-essentiality map^39^, single-cell co-expression gene pairs^42^, and two compendia of physical protein-protein contact maps^40,41^ to comprise the positive testing data. We filtered the testing set to remove any pairs that occur in the training set to avoid data leakage.

We trained a series of machine learning models using a temporal split (described above). Co-occurring genes were used as positive training examples and negative training examples were randomly sampled gene pairs extensively filtered to avoid accidental positives: we ensured that negative pairs did not overlap our training or test sets, did not share a Gene Ontology^19^ process, did not exist in the STRING database^43^, and were separated by at least 100,000 base pairs on the hg38 reference genome. Other training details mirror previously described methods.

