## Supplementary Information for "Domain-specific embeddings uncover latent genetics knowledge"

**S. S. Ho[1], R. E. Mills[1,2]**

[1]Department of Human Genetics, University of Michigan, Ann Arbor, MI, USA

[2]Department of Computational Medicine and Bioinformatics, University of Michigan, Ann Arbor, MI, USA

**Supplementary Fig. 1**: Expression prediction using transformed-based embeddings

**Supplementary Fig. 2**: Precision-recall curves comparing gene-disease association prediction performance with different cut-off years

**Supplementary Fig. 3**: Decreasing size of gene-disease association test-set over time.

**Supplementary Fig. 4**: Precision-recall curves comparing cancer driver gene prediction performance with different cut-off years.

**Supplementary Fig. 5**: Relationship between cancer driver gene prediction performance and test set size

**Supplementary Fig. 6**: Precision-recall curves for temporally-split interaction prediction models.

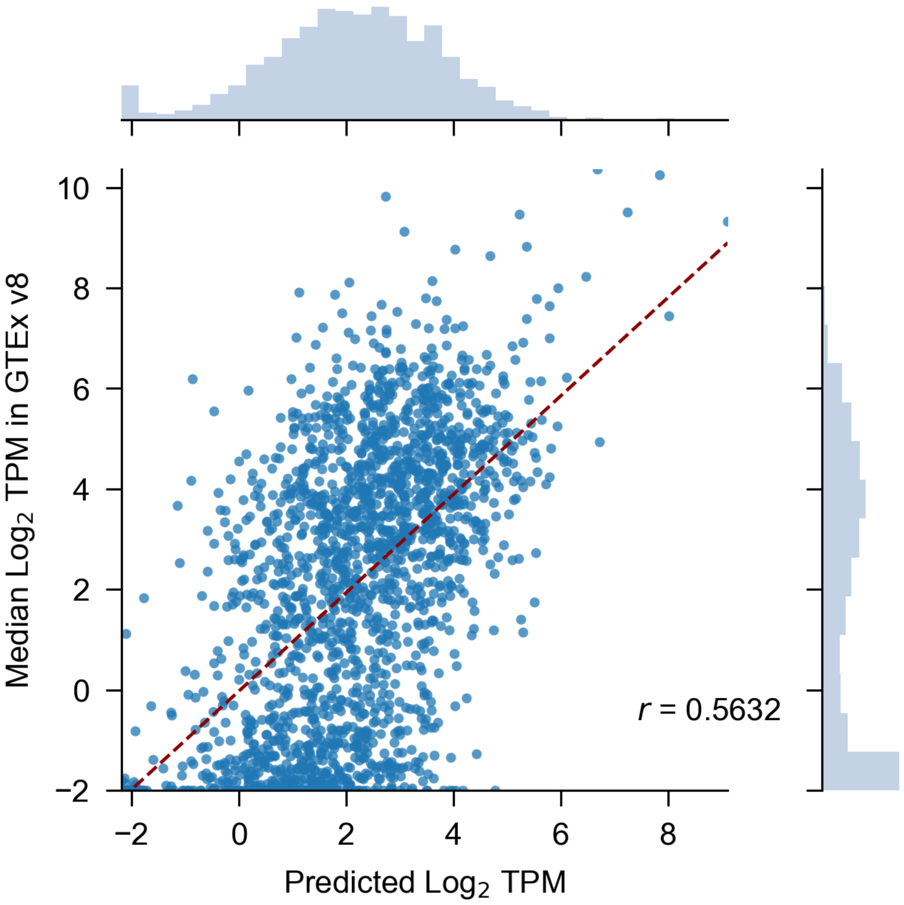


**Supplementary Fig. 1 | Expression prediction using transformed-based embeddings. a**, Scatter plot of predicted versus expected gene expression hold-out test genes (chr 1) using embeddings derived from GenePT.

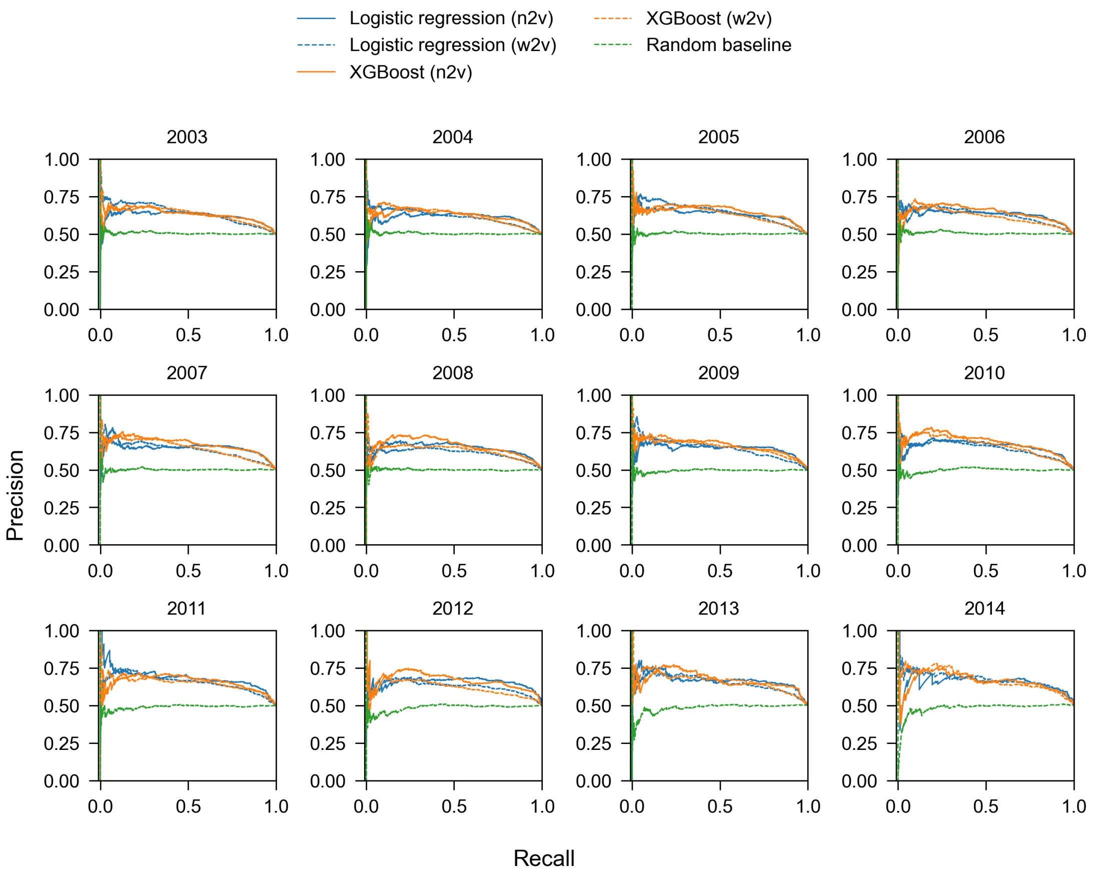


**Supplementary Fig. 2 | Precision-recall curves comparing gene-disease association prediction performance with different cut-off years.** Precision-recall curves on the test set stratified by year cut-off.

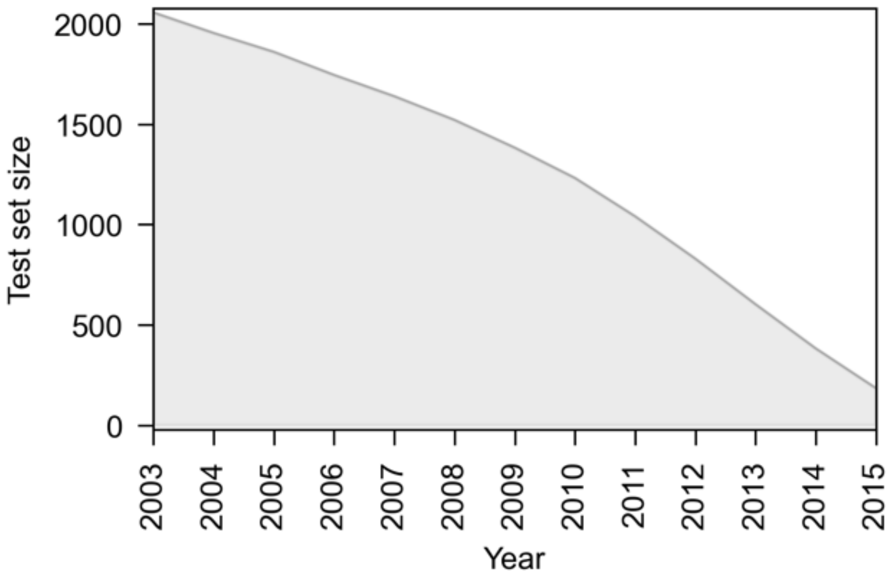


**Supplementary Fig. 3 | Decreasing size of gene-disease association test-set over time.**

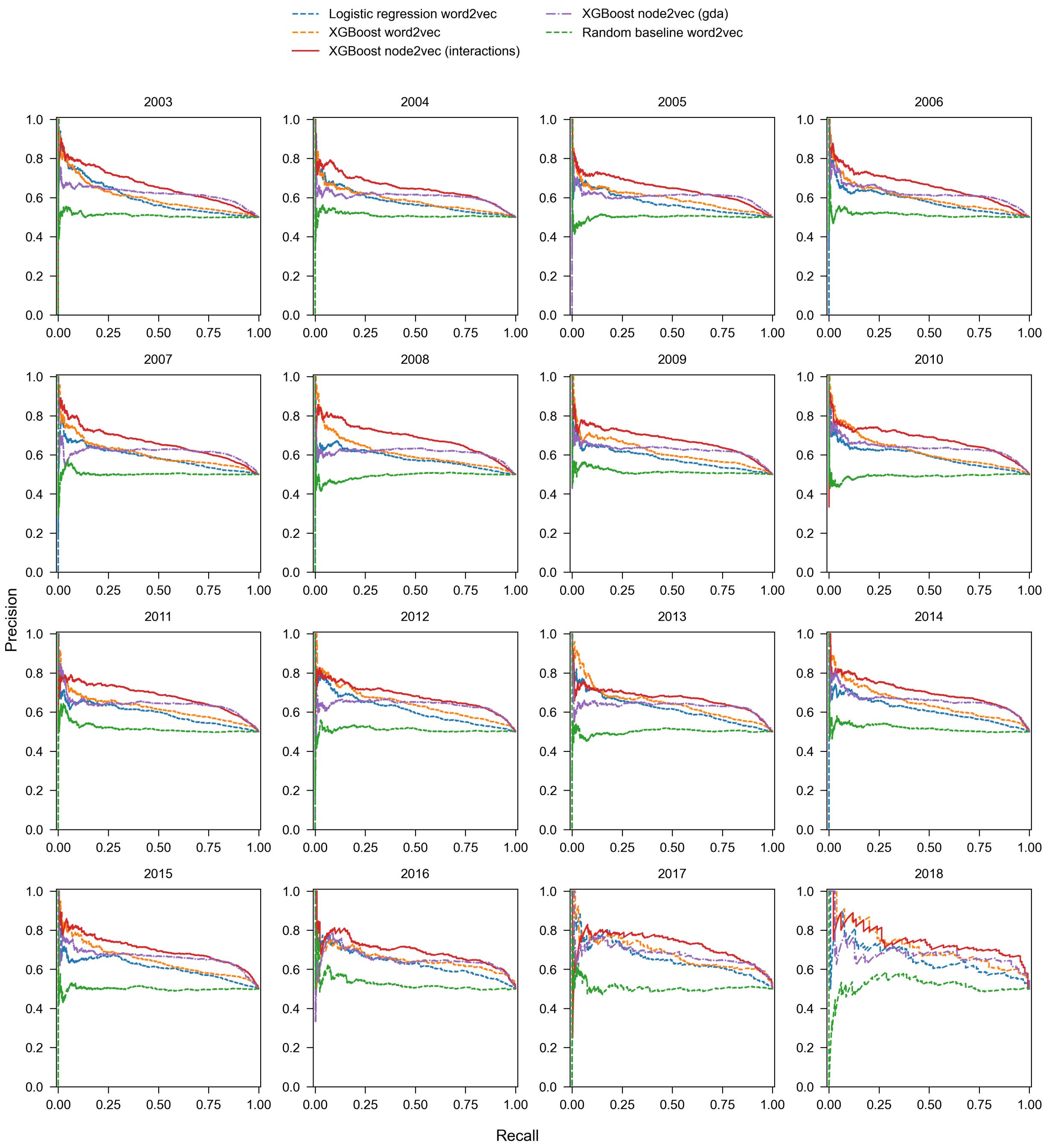


**Supplementary Fig. 4 | Precision-recall curves comparing cancer driver gene prediction performance with different cut-off years.** Panels represent performance for specific years (2003-2018). XGBoost model leveraging node2vec gene-gene co-occurrence data (red) consistent demonstrates superior performance across the temporal splits.

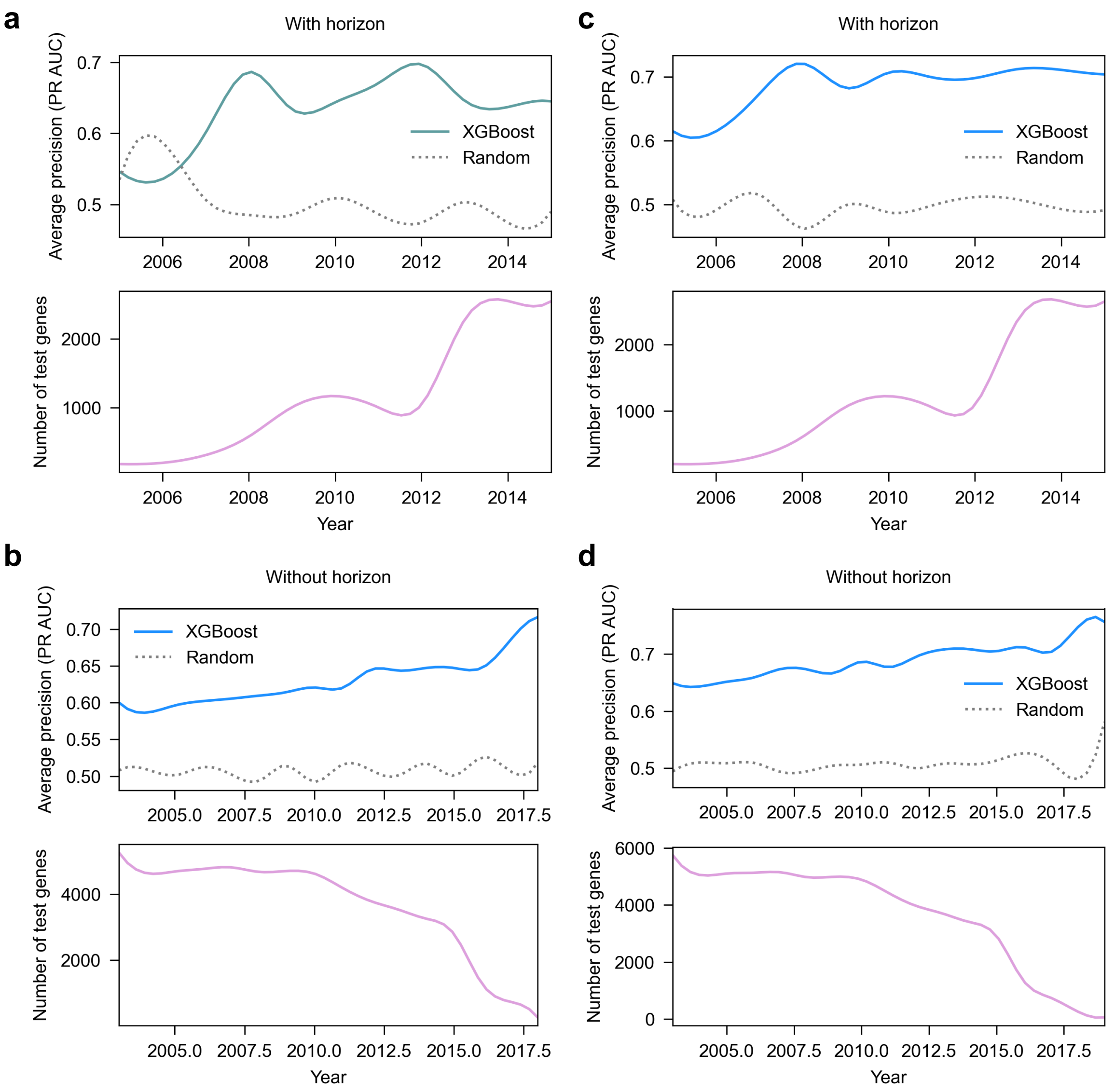


**Supplementary Fig. 5 | Relationship between cancer driver gene prediction performance and test set size.** XGBoost models trained on word2Vec embeddings (**a**,**b**) and node2vec embeddings (**c**, **d**) with corpora from different temporal cutoffs. Each model is evaluated on cancer-associated genes discovered after its training cutoff year. Top panels show precision-recall AUC values compared to a random baseline; bottom panels show the number of test genes for each year. "With horizon" (**a**,**c**) indicates testing on all future cancer genes, while "Without horizon" (**b**,**d**) restricts testing to genes discovered within three years of the training cutoff.

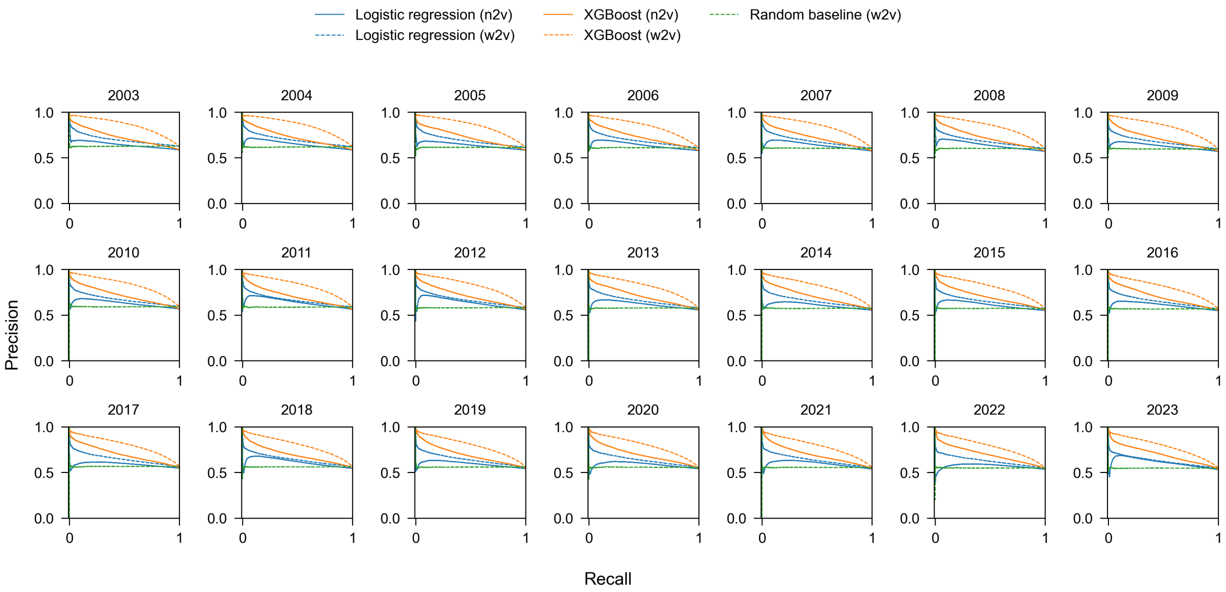


**Supplementary Fig. 6 | Precision-recall curves for temporally-split interaction prediction models.** Panels display precision-recall curves for multiple interaction prediction models trained on corpora up to the specified year. Logistic regression and gradient-boosted models were trained using embeddings from word2vec and node2vec and compared against a random baseline (green). Word2vec models consistently outperform node2vec models and exhibit stable performance across years.
